# Reconstructing phylogenies of metastatic cancers

**DOI:** 10.1101/048157

**Authors:** Johannes G. Reiter, Alvin P. Makohon-Moore, Jeffrey M. Gerold, Ivana Bozic, Krishnendu Chatterjee, Christine A. Iacobuzio-Donahue, Bert Vogelstein, Martin A. Nowak

## Abstract

Reconstructing the evolutionary history of metastases is critical for understanding their basic biological principles and has profound clinical implications^1–3^. Genome-wide sequencing data has enabled modern phylogenomic methods to accurately dissect subclones and their phylogenies from noisy and impure bulk tumor samples at unprecedented depth^4–7^. However, existing methods are not designed to infer metastatic seeding patterns. We have developed a tool, called Treeomics, that utilizes Bayesian inference and Integer Linear Programming to reconstruct the phylogeny of metastases. *Treeomics* allowed us to infer comprehensive seeding patterns for pancreatic^8^, ovarian^9^, and prostate cancers^10,11^. Moreover, Treeomics correctly disambiguated true seeding patterns from sequencing artifacts; 7% of variants were misclassified by conventional statistical methods. These artifacts can skew phylogenies by creating illusory tumor heterogeneity among distinct samples. Last, we performed *in silico* benchmarking on simulated tumor phylogenies across a wide range of sample purities (30-90%) and sequencing depths (50-800x) to demonstrate the high accuracy of Treeomics compared to existing methods.

---

Genetic evolution underlies our current understanding of cancer^12–14^ and the development of resistance to therapies^15,16^. The principles governing this evolution are still an active area of research, particularly for metastasis, the final biological stage of cancer that is responsible for the vast majority of deaths from the disease. Although many insights into the nature of metastasis have emerged^2,17,18^, we do not yet know how malignant tumors evolve the potential to metastasize, nor do we know the temporal or spatial rules governing the seeding of metastases at sites distant from the primary tumor^1,3,19–21^.

In order to better understand the process of metastasis, researchers have reconstructed the temporal evolution of patients’ cancers from genome sequencing data^22–25^. Thus far, phylogenomic analysis has largely focused on the subclonal composition and branching patterns of primary tumors^11,26,27^. The evolutionary relationships *among* metastases are equally important but have less often been determined for several reasons^9,10,28^. First, comprehensive data sets of samples from spatially-distinct metastases in different organs are rarely available. Second, most advanced cancer samples are derived from patients who have been treated with toxic and mutagenic chemotherapies, imposing a variety of unknown constraints on genetic evolution and its interpretation. Third, tumors are composed of varying proportions of neoplastic and non-neoplastic cells, and inferring meaningful evolutionary patterns from such impure samples is challenging^29,30^. Moreover, the situation for solid tumors differs from that of “liquid tumors”, where mutant allele fractions are high and can be easily determined from cytological analysis. Fourth, chromosome-level changes, including losses, are frequently observed in cancers, and previously acquired variants can be lost (i.e., some variants are not “persistent”). Fifth, even when performed at high depth, next-generation sequencing coverage is always non-uniform, resulting in different amounts of uncertainty at different loci within the same DNA sample as well as among different samples at the same locus. Finally, evolutionarily informative genetic differences among the founding cells of distant metastases tend to be rare^8^.

The variety of methods that have recently been used to infer evolutionary relationships among tumors underscore these complicating factors and the need for a robust phylogenomic approach. The methods include those based on genetic distance^9^, maximum parsimony^27,28,31^ clonal ordering^14,25,32^ and variant allele frequency^33–35^. Modern phylogenomic methods classify variants based on the observed variant allele frequencies (VAFs), account for varying ploidy and neoplastic cell content, and reconstruct comprehensive phylogenies^4–7, 36–38^. However, as we will show below, in the case of reconstructing the evolution of metastases, these methods suffer from the low number of informative variants and may fail to identify the subclones that gave rise to the observed seeding patterns. Classical phylogenetics assumes that the individual traits are known with certainty^29^. Consequently, these methods struggle with noisy high-throughput DNA sequencing data and do not exploit the full potential of these data due to the error-prone binary present/absent classification of variants. Furthermore, many of the methods used for inferring cancer evolutionary trees are based on those designed for more complex evolutionary processes involving sex and recombination^20^.

## RESULTS

### Evolutionarily incompatible mutation patterns

To illustrate our approach, we first focused on the data of a treatment-naïve pancreatic cancer patient Pam03^8^ (Fig. 1). WGS (whole-genome sequencing; coverage: median 51x, mean 56x) as well as deep targeted sequencing (coverage: median 296x, mean 644x) was performed on ten spatially-distinct samples: two from the primary tumor and eight from distinct liver and lung metastases (Online Methods and ref. 8). Estimated purities ranged from 18% to 46% per sample (Fig. S2; Supplementary Information), typical for low-cellularity cancers (Fig. 1). Founder variants (present in all samples) and unique variants (present in exactly one sample) are parsimony-uninformative and hence irrelevant for the branching in an evolutionary tree. Parsimony-informative variants (variants present in some but not in all samples) exhibited contradicting mutation patterns when we tried to reconstruct a phylogeny consistent with the evolutionary processes underlying tumor progression using conventional methods. Identifying the evolutionarily compatible variants is known as the “binary maximum compatibility problem” and has been widely studied for decades^39–43^. A strict binary present/absent classification can be very problematic due to the wide coverage distribution in sequencing data^43^. For example, likely clonal variants in the driver genes *ATM* and *KRAS* would be classified as absent in sample LuM 2 because both were covered only fourteen times and were mutated only once (Fig. 1b; Supplementary Table S2). We developed a Bayesian inference model to determine the posterior probability of whether a variant was or was not found in each sequenced lesion rather than rely on a binary input (“present” or “absent”; Fig. 1b; Online Methods). This generalization, formalized as a Mixed Integer Linear Program^44^ (MILP), enabled us to simultaneously predict sequencing artifacts and infer phylogenies in a remarkably robust fashion.

**Figure 1:**
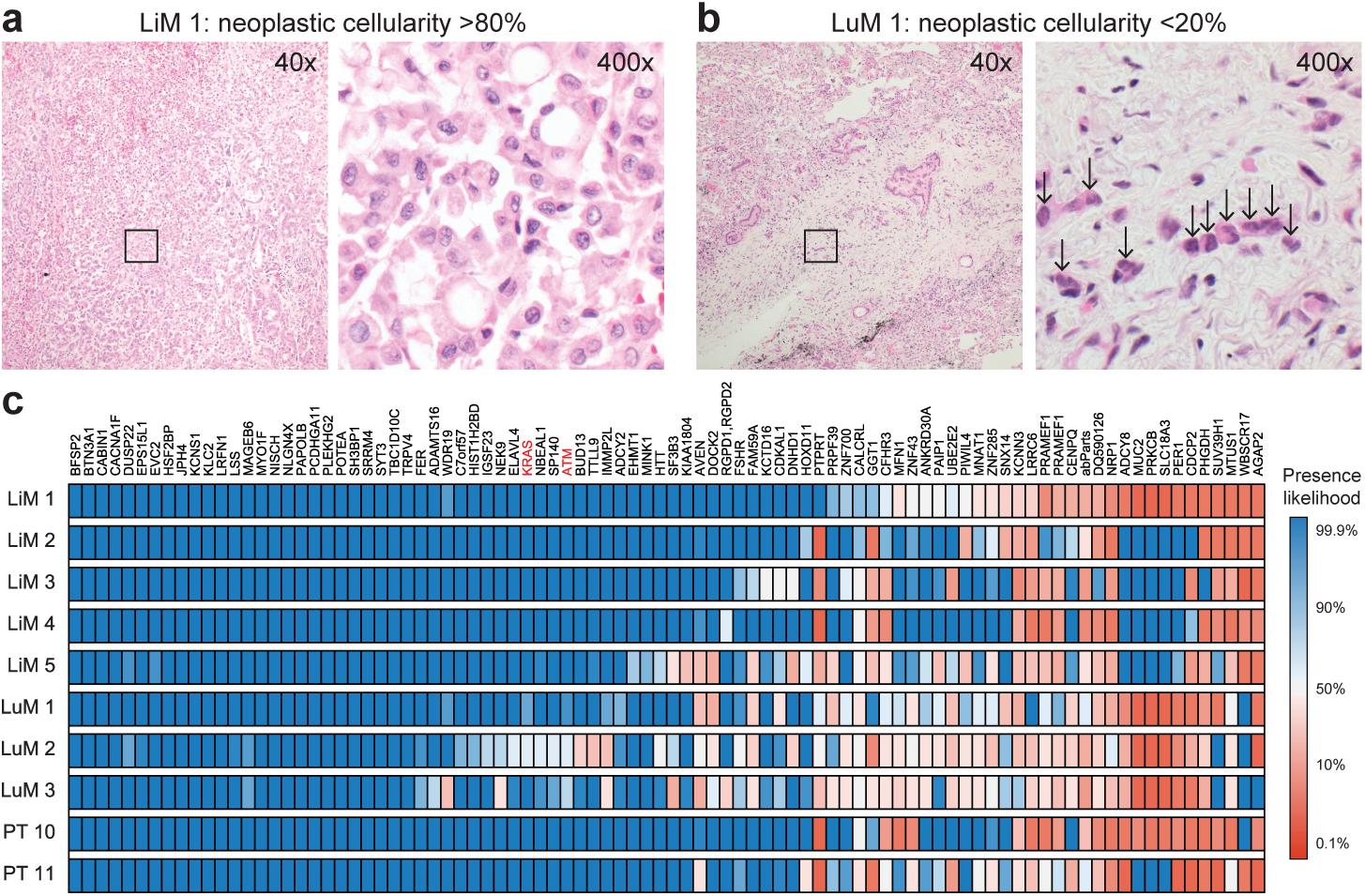
Tumor heterogeneity across lesions of pancreatic cancer patient Pam03. a, b. | Histology at low (40×) and high (400×) power of liver metastasis LiM 1 and lung metastasis LuM 1, with estimates of neoplastic cellularity determined by pathological review. Arrows highlight the few cancer cells in LuM 1. **c** | Heatmap depicting the posterior probability (p) that a variant is considered as present in deep targeted sequencing data. Top five rows show samples from five distinct liver metastases (LiM 1-5); the following three rows show samples from three distinct lung metastases (LuM 1-3); the bottom two rows show different parts of the primary tumor (PT 10-11). Dark blue corresponds to a variant being present with probability > 99.9% and dark red corresponds to being absent with probability > 99.9%. In some samples the mutation status for the most likely clonal driver mutations in *ATM* and *KRAS* is unknown.

Two variants are evolutionary compatible if there exists an evolutionary tree where each variant is only acquired once and never lost. This condition is known as the perfect (the same variant is not independently acquired twice; infinite sites model^45^) and persistent (acquired variants are not lost; no back mutation) phylogeny assumption – the basic principle of modern tumor phylogeny reconstruction methods^4–7,36^. In our case the mutation pattern of a variant is given by the set of samples where the variant is present (Fig. S1). Therefore, two somatic variants *α* and *β* are evolutionarily incompatible if and only if samples with the following three patterns exist: (i) variant *α* is absent and *β* is present, (ii) *α* is present and *β* is absent, and (iii) both variants are present. Because somatic variants are by definition absent in the germline, *α* and *β* are evolutionarily incompatible and no perfect and persistent phylogeny can explain these data (Fig. S1). As expected, based on conventional binary present/absent classification of variants, a perfect and persistent tree consistent with the observed (noisy) data of Pam03 cannot be inferred. We show that such a phylogeny indeed exists but that it is hidden behind misleading technical and biological artifacts, mostly resulting from insufficient coverage or low neoplastic cell content (Fig. S2).

### Identifying evolutionarily compatible mutation patterns

To account for inconclusive data, we utilize a Bayesian inference model to calculate the probability that a variant is present in a sample (Fig. 1c, Online Methods). Using these probabilities for each individual variant, we calculated reliability scores combining the evidence for each possible mutation pattern across all variants. We constructed an evolutionary conflict graph where the nodes were determined through analysis of all mutation patterns. Each node was assigned a weight provided by the calculated reliability scores (Fig. S3). If two nodes (mutation patterns) were evolutionarily incompatible, an edge between the corresponding nodes was added. We aimed to identify the set of nodes that maximized the sum of the weights (reliability scores) when no pair of nodes was evolutionarily incompatible. This maximal set represents the most reliable and evolutionarily compatible mutation patterns (Supplementary Information). To evaluate the confidence in the identified evolutionarily compatible mutation patterns, we performed bootstrapping on the provided variants.

### Predicting putative artifacts in sequencing data

The solution obtained with the MILP directly provided the most likely evolutionarily compatible mutation pattern for each variant. By comparing our inferred classifications to conventional binary classifications, Treeomics predicted putative sequencing or biological artifacts in the data (Fig. 2a,b). The conventional classifications differed in 9.2% of the variants in Pam03 (83 putative artifacts from 90 variants across 10 samples; Fig. 2b). As expected, the majority (77) of the differences were caused by putative false-negatives in the binary classification that were inferred to be present by Treeomics. Sixty-four of these putative false-negatives had relatively low coverage (median sequencing depth: 18), explaining how they could easily be misclassified as absent given the low neoplastic cell content in these samples. Accordingly, many of these under-powered false-negatives occurred in samples with the lowest coverage (liver metastasis LiM 5, lung metastases LuM 2-3) or lowest neoplastic cell content (LuM 1; Fig. S2). In LuM 2, the driver gene mutation *KRAS* was incorrectly classified as absent by conventional means though it is most likely a clonal founding mutation and was present at a VAF of 19% in the original WGS sample (Supplementary Table S1). Similarly the driver gene mutation *ATM* was incorrectly classified as absent in two samples (VAF 18% and 19% in the WGS data). Although manual review of these samples revealed mutant reads in *KRAS*, it is not scalable to manually review every putative variant detected by next generation sequencing. Some variants contained false-negatives across many samples, indicating that these variants were generally difficult to call. Remarkably, 91% (58/64) of the predicted under-powered false-negatives were either significantly present in the WGS data (45/58; mostly at higher coverage than in the targeted sequencing data), or the genomic region of the variant possessed a low alignability score^46,47^ (13/58; Supplementary Table S1).

**Figure 2:**
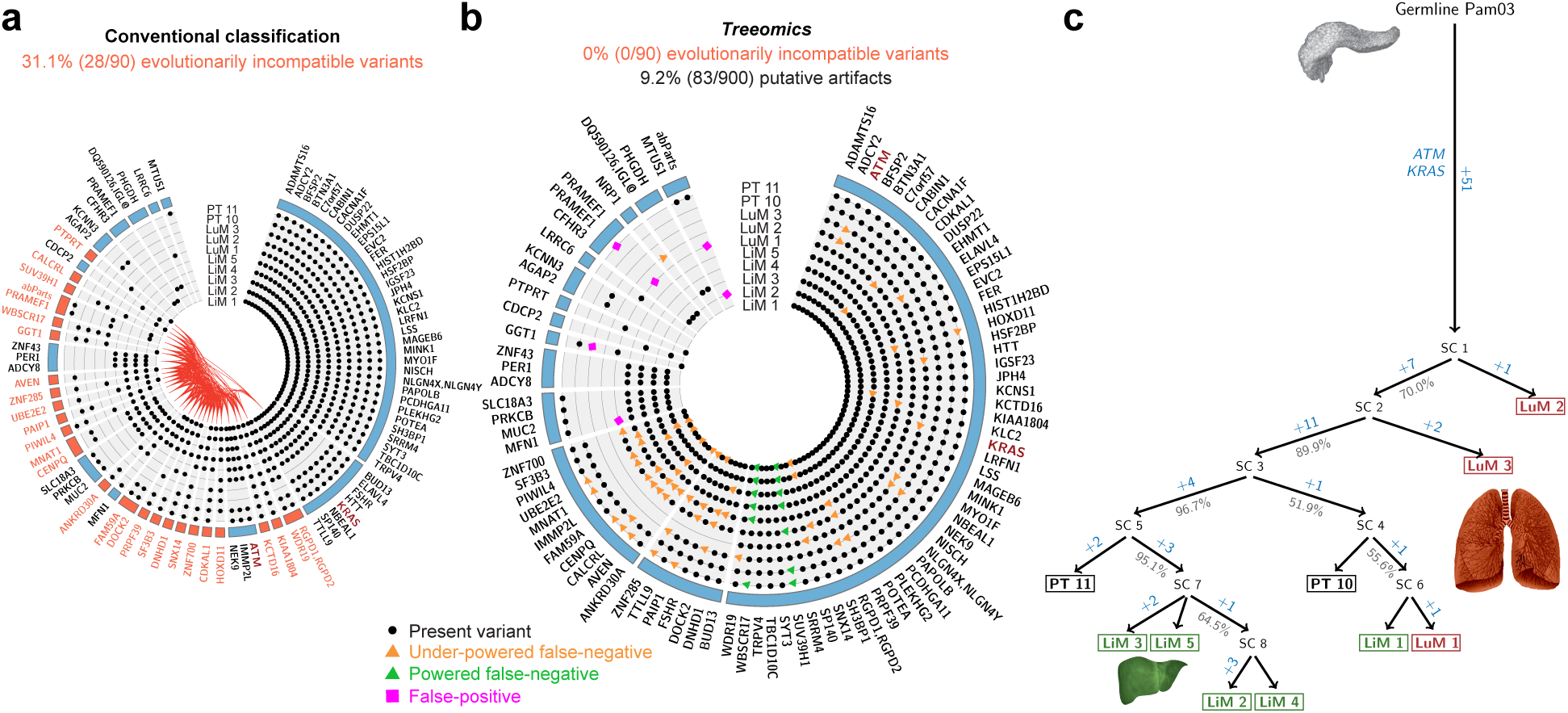
Treeomics simultaneously identified putative artifacts and inferred the evolutionary history of Pam03. a, b. | Variants shown in Fig. 1c are organized as evolutionarily-defined groups (nodes). Blue colored nodes are evolutionarily compatible and red colored nodes are evolutionarily incompatible. Based on conventional present/absent classification, 31.1% of the variants were evolutionarily incompatible (**a**). The incompatibilities are demarcated by red lines (edges) in the center of the circle that connect each pair of incompatible nodes. Based on a Bayesian inference model and an Integer Linear Program, Treeomics identified the most likely evolutionarily compatible mutation pattern for each variant (**b**; Online Methods). This method predicted that 9.2% (83/900) of the variants across all samples were misclassified and thereby caused the evolutionary incompatibilities shown in panel **a**. 76% of the predicted artifacts were validated in the WGS data, among those were artifacts in ATM and *KRAS.* **c** | Reconstructed phylogeny from the identified evolutionarily compatible mutation patterns in panel **b**. Gray percentages indicate bootstrapping values from 1000 samples. SC indicate predicted subclones. Lung metastases (LuM 1-3) are depicted in red; Liver metastases (LiM 1-5) are depicted in green; Primary tumor samples (PT 10-11) are depicted in black.

For two variants sequenced at high depth, Treeomics predicted 13 putative false-negatives. The WGS data confirmed sequencing artifacts in these two variants but indicated that 4 likely false-positives (all absent in the WGS data) induced Treeomics to predict 13 false-negatives rather than 4 false-positives (Supplementary Table S1). Of the 6 putative false-positives (pink squares in Fig. 2b), 83% (5/6) were classified as absent in the original WGS data and their median VAF was 1.3% (Supplementary Table S1). In total, 76% (58 putative false-negatives + 5 putative false-positives; 63/83) of the predicted artifacts were successfully validated. Hence, we verified that at least 7% (63/900) of the variants were misclassified by conventional binary classification. If a phylogenomic method does not account for sequencing artifacts, the mutation patterns of a large fraction of variants will often be inconsistent with any inferred evolutionary tree. In Pam03, the mutation patterns of 31.1% (28/90) of the variants would be evolutionarily incompatible (Fig. 2a). These putative artifacts may also help to explain the observed high tumor heterogeneity in earlier studies and the recently reported tumor similarity when sequencing depth is increased^8,27^.

### Inferring evolutionary trees

From the identified mutation patterns, Treeomics inferred an evolutionary tree rooted at the germline DNA sequence of the pancreatic cancer patient Pam03 (Fig. 2c). We found strong support for an evolutionarily related group of geographically distinct lesions: samples LiM 2-5 (liver metastases) and PT 11 (primary tumor). These results suggest that a recent parental clone of PT 11 seeded these liver metastases. We also found the same evolutionary relationship by using the low-coverage WGS data (Fig. S4). In contrast to the targeted sequencing data, the WGS data indicated that lung metastasis LuM 1 was more closely related to LuM 2 and LuM 3. Though the low neoplastic cell content prevents a definite conclusion about the seeding subclone of LuM 1, the reconstructed phylogeny strongly suggests that the liver metastasis LiM 1 was seeded from a genetically different subclone than all other liver metastases. This diversity in seeding subclones was also found in another treatment-naïve pancreatic cancer patient (Pam01) whose data similarly indicated that liver metastases were seeded from genetically distinct subclones (Fig. S5). The phylogeny of Pam01 suggested that distinct subclones of the primary tumor gave rise to not just different liver metastases but also different lymph node metastases. This observation suggests that subclones are not necessarily predisposed to seeding at a particular site. In contrast, the phylogeny of Pam02 revealed that all liver metastases except one (LiM 7 with low median coverage: 28) were very closely related to each other and to various regions of the primary tumor – indicating recent divergence (Fig. S6). Pam02’s pancreatic cancer might have been expanded very rapidly with only 0.5 months from diagnosis to death compared to 7 and 10 months for Pam01 and Pam03.

To further validate our approach, we reanalyzed data from high-grade serous ovarian cancers^9^. We were able to reproduce all phylogenetic trees of Bashashati et al.^9^ except for Case 5. In this case, the authors reported an early divergence of sample 5c while Treeomics suggested a later divergence (Fig. S7c). Comprehensive analysis of their data (reinterpreted in Fig. S7a,b) revealed that their tree either required that several variants (including two driver gene mutations and multiple indels) occurred independently twice or that two mutations in the driver genes *ABL1* and *MDM4* were lost. Both possibilities seem unlikely (Fig. S7 and Fig. 1D in ref. 9); this discrepancy was also identified by Popic et al. (ref. 6). Treeomics did not require these implausible scenarios to construct an otherwise similar tree. Distance-based methods, such as those used by Bashashati et al., can be compromised by large differences in the number of acquired mutations among samples; sample 5c had twice as many mutations than most other samples.

We also reanalyzed a comprehensive data set from prostate cancers^10^. Treeomics confirmed the majority of results but also further refined others. For example, for patient A32, Gundem et al. (2015) reported an inconclusive evolutionary tree due to evolutionary incompatible subclones present at low frequencies. Our method used the strong evidence for mutation patterns C, E and D, F (see Extended Data Figure 3p,q in ref. 10) and was thereby able to illuminate the evolutionary relationships among these samples in a conclusive fashion (Fig. S8).

If multiple subclones with spatially-distinct evolutionary histories (e.g., metastasis reseeding) were present in the same sample at detectable frequencies, conventional phylogenetic approaches would be unable to separate their evolutionary trajectories. In these scenarios, evolutionarily incompatible mutation patterns with high reliability scores were identified and utilized to detect these subclones and to infer separate evolutionary histories (Fig. S9b; Online Methods; Supplementary Information). For both the prostate cancer data of case A22^10^ (Fig. S9) and of case 6^11^ (Fig. S10), Treeomics identified subclonal structures and separated their evolutionary trajectories without requiring high purity samples or deep sequencing data.

### *In silico* benchmarking demonstrates high accuracy

We implemented a stochastic continuous-time multi-type branching process to imitate the genetics of distinct metastases seeded according to an evolving cancer^21,48^ (Fig. 3; Online Methods). Based on the simulated ground truth data, we compared the performance of Treeomics with conventional phylogenetic methods (Maximum Parsimony and Neighbor Joining) and a modern phylogenomic method (PhyloWGS^7^) across sample purities of 30% to 90% and sequencing depths of 50 to 800 (Fig. 3c-f, Figs. S11-S12) representing the range of typical sample quality. We investigated an unprecedented total of 37,500 independently simulated phylogenies comprised of 75 different combinations of sample purity, mean sequencing depth and intermetastatic mixing. All simulated phylogenies can be downloaded from a github repository (Online Methods). The benchmarking results demonstrate that phylogenies obtained from low coverage WES data show high error rates independent of the used method. Even for high purities, one third of the inferred branchings tend to be wrong. For mean coverages of 100 and above, the error rates drop dramatically and phylogenies can be accurately reconstructed. For a mean coverage of 200 and a neoplastic cell content of 75%, Maximum Parsimony makes about twice the number of errors compared to Treeomics (Fig. 3d, Fig. S11).

**Figure 3:**
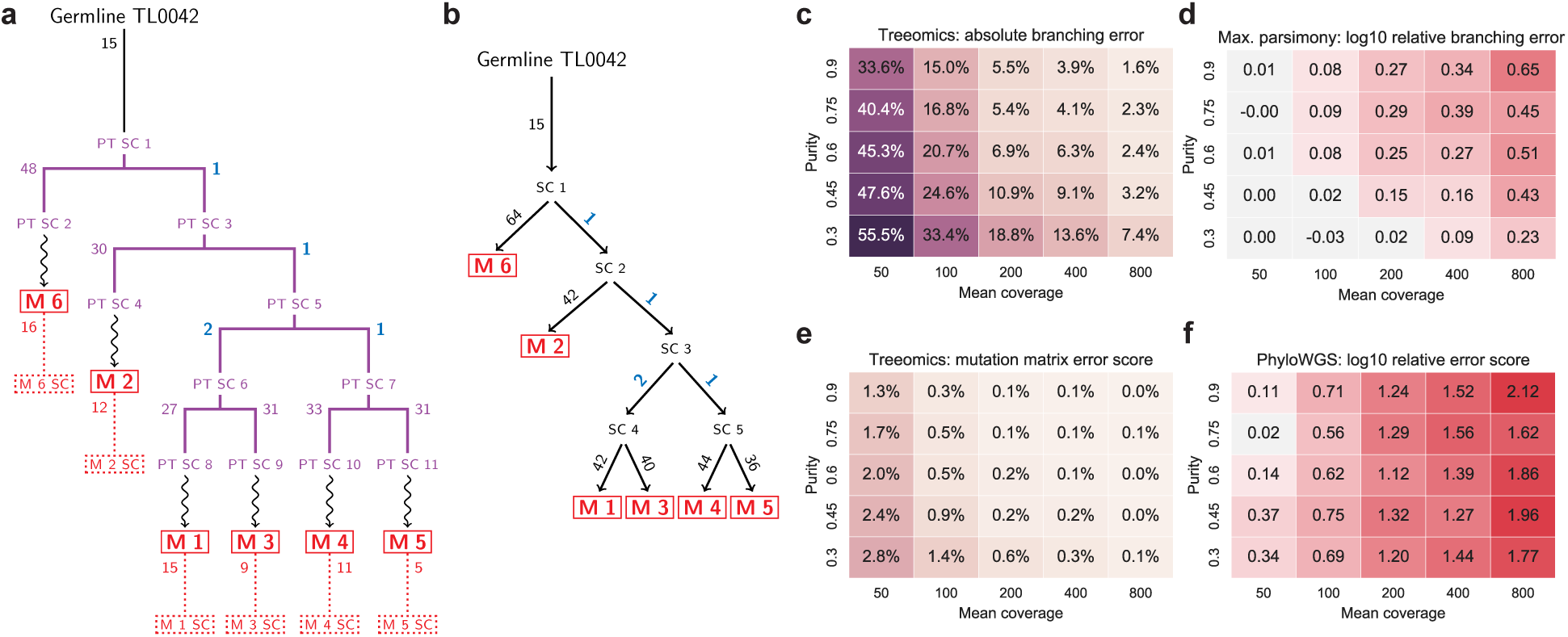
In silico benchmarking demonstrates high accuracy of Treeomics across varying sample purity and mean sequencing depth compared to existing methods. a. | Simulated metastatic progression according to a stochastic branching process^20,47^. Metastases (M 1-6) are numbered in chronological order of their seeding. Purple lines indicate evolution along lineages within the primary tumor. Blue numbers correspond to the parsimony informative variants. Numbers in red denote subclonal variants acquired after the seeding of the metastasis. SC indicates subclone. Dotted boxes illustrate biopsies. **b** | Treeomics correctly reconstructed the simulated phylogeny in panel **a** by identifying the few parsimony informative variants (blue). Private mutations acquired in the primary tumor are indistinguishable from subsequently acquired mutations. **c-f** j Benchmarking across 12,500 simulated phylogenies with six metastases. Branching error and mutation matrix error score dropped drastically with increasing sequencing depth. For purities above 60% and coverages above 200, Maximum parsimony made between 80% and 340% more errors than Treeomics. PhyloWGS exhibited error scores more than 10-fold higher than those of Treeomics in most considered scenarios.

Current subclone inference algorithms do not directly reconstruct phylogenies of distinct sites as Treeomics does but infer joint phylogenies of variants, which are sometimes simultaneously grouped into subclones^6,7,36–38^. To enable a comparison of these slightly different methodologies, we developed a mutation matrix error score (similar as in ref. 6) that checks (i) if variants of the same subclone were indeed assigned to the same subclone and (ii) if the ancestral relationship among variants was correctly determined (Online Methods). Since the runtime of PhyloWGS increases significantly with the number of variants, we removed all noisy, private variants in the input for PhyloWGS. Despite this advantage by removing potential noise, Treeomics outperformed PhyloWGS in all considered scenarios (Fig. 3f). In the majority of scenarios, the error score of PhyloWGS was more than 10-fold higher than the error score of Treeomics. We also note that the runtime of PhyloWGS was around 5-10 hours per simulated phylogeny (in total ~200,000 core computing hours), while Treeomics needed a few seconds per case (in total ~50 core computing hours).

## DISCUSSION

The new approach described here efficiently reconstructs the evolutionary history, detects potential artifacts in noisy sequencing data, and finds subclones of distinct origin. The evolutionary theory of asexually evolving populations combined with Bayesian inference and Integer Linear Programming enabled us to infer detailed phylogenomic trees with significantly fewer errors than existing methods. In contrast to other tools, Treeomics accounts for putative artifacts in sequencing data and can thereby infer the branches where somatic variants were acquired as well as where some may have been lost during evolution, presumably through losses of heterozygosity resulting from chromosomal instability^49^. The branching in the inferred trees sheds new light on the seeding patterns of particular metastatic lesions^1,20^. For example, in contrast to colon cancer, where liver metastases are assumed to seed lung metastases^50^, our results suggest that this may not be the case in pancreatic cancer. The reconstructed phylogenies also indicate that distinct subclones in the primary tumor were equally capable to seed metastases in the same and in different organs (Fig. S5). The evolutionary rules of natural metastatic cancers leading to the highly non-random pattern of metastases in Pam03 are just beginning to emerge.

Despite these detailed reconstructed phylogenies, there are several limitations that should not be neglected. Without additional data, even completely correct cancer phylogenies do neither directly provide information about the temporal ordering in which metastases were seeded nor about the anatomic location of the seeding subclones. For example, metastasis M6 was seeded last but diverged first in the simulated phylogeny (Fig. 3a). Furthermore, a single seeding event cannot be distinguished from multiple seeding events from the topology of the reconstructed tree alone (see ref. 20). Incorporating multiple samples of the primary tumor and inferring the acquired mutations on the individual branches can provide evidence about the location of the seeding subclone and the timing of the seeding event. For example, the genetic similarity of the primary tumor sample PT 11 and the liver metastases LiM 2-5 suggests multiple seeding events from a recent ancestor of PT 11. Moreover, phylogenomic approaches could incorporate estimated growth rates and mutation rates to better quantify the probability of metastasis-to-metastasis spread.

We have designed Treeomics from first principles to directly handle ambiguity in high-throughput sequencing data, including samples with low neoplastic cell content or coverage. The mutation patterns and their evolutionary conflict graph form a robust data structure and consequently the painful task of semi-automatic filtering becomes unnecessary. As a result of the Bayesian confidence estimates for the individual variants, this method can infer more robust results than traditional phylogenetic methods, which employ a binary representation of sequencing data (Fig. 2a). Furthermore, as shown above, distance-based methods can produce results inconsistent with the evolutionary theory of cancer as they often ignore knowledge of biological phenomena specific to neoplasia (Fig. S7). We note that PhyloWGS and other subclone inference methods have not been designed to reconstruct phylogenies based on these few genetic variants that determine the evolutionary history of metastases. The key difference between these approaches is that Treeomics assumes that mixing of subclones from two spatially-distinct sites is rare. Treeomics therefore works extremely well among metastases, however would not be applicable for liquid cancers. On the contrary, tools like PhyloWGS work extremely well in liquid cancers. Last, we compared our results to AncesTree^36^, which roughly identified the evolutionarily related samples in Pam03 but excluded 70% (63/90) of the variants (among them the driver gene mutations in *KRAS* and *ATM)* in the inferred phylogeny due to evolutionary incompatibilities (Fig. S13).

At present, Treeomics only employs nucleotide substitutions and short insertions and deletions – a subset of the available information. The benchmarking results demonstrate that a single mutation varying in two samples is typically sufficient for Treeomics to infer the correct evolutionary history (Fig. 3); a crucial property given the high genetic similarity of metastases^8^. Other types of data, such as copy number alterations, structural variations and DNA methylation, could be incorporated into Treeomics to further improve the accuracy of the inferred results^51–54^.

Day and Sankoff showed that inferring the most likely evolutionary trajectories is a computationally challenging problem (*NP*-complete^39^). Sophisticated approximation algorithms have been developed in the context of language and cancer evolution^40,42,43^. However, medium-sized instances of *NP*-complete problems are no longer intractable due to the enormous engineering and research effort that has been devoted to ILP solvers. The MILP^44^ formulation enables an efficient and robust analysis of large datasets. We prove that an approximation algorithm that would guarantee that its solution is at most 36.06% worse than the optimal solution cannot exist unless the complexity class *P*=*NP* (Supplementary Information, Theorem 1). Salari et al.^43^ explored a related approach but approximated two *NP*-complete problems, possibly leading to suboptimal results. Treeomics produces a mathematically guaranteed to be optimal result without convergence or termination issues. Note that a mathematical optimal solution is not necessarily equivalent to the biological truth, specifically in the case of low neoplastic cell content or coverage (Fig. 3c,e). MILPs may also be useful in other areas of phylogenetic inference where methods with strong biological assumptions (e.g. constant mutation rates or specific substitution profiles) are not applicable or are computationally too expensive to obtain guaranteed optimal solutions.

## ONLINE METHODS

### DNA sequencing design and validation

As described in detail in ref. 8, sequencing data were generated in two stages. First, genomic DNA from 26 tumor samples (20 metastases and 6 primary tumor sections) was evaluated by 60x whole genome sequencing (WGS) using an Illumina Hi-Seq 2000. Importantly, genomic DNA from the normal tissue of each patient was used to facilitate identification of somatic variants. We obtained an average coverage of 69x with 97.5% of bases covered at >10x, revealing a total of 127,597 putative coding and noncoding somatic mutations, (average of 4,908 per sample). To limit the artifacts generated by WGS and alignment, we filtered the putative variants using several quality parameters, including read directionality, mutant allele frequency detected in the normal, known human SNPs, and the number of independent tags at each site.

Second, we utilized a targeted sequencing approach to independently screen every mutation that we observed to be of high quality in at least one WGS tumor sample. Briefly, probes for capture were designed to flank each potential mutant base (n = 2105) and libraries were prepared for the original 26 WGS samples. Using an Illumina chip-based approach, we successfully aligned, processed, and validated 381 mutations (range 106-164 per patient) at an average sequencing depth of 731x (Supplementary Tables S2, S3, S4). In addition to the increased coverage and sensitivity of targeted sequencing, both sequencing approaches generated independent datasets in which we could directly compare putative variants *in silico* among many tumors within a patient. Additional details regarding patient selection, processing of tissue samples and DNA extraction and quantification can be found in ref. 8.

### Bayesian inference model

To compute reliability scores for each mutation pattern, we extract posterior probabilities for the presence and absence of a variant in a sample from a Bayesian binomial likelihood model of error-prone sequencing. If *f* is the true fraction of variant reads in the sample, *π* is our prior belief about *f*, and *e* is the sequencing error rate, the posterior distribution *P* of *f* given *N* total reads and *K* variant reads is

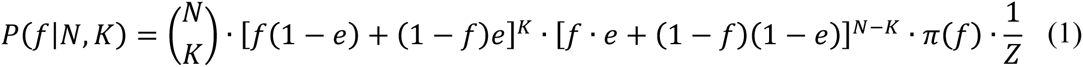

where *Z* is a normalizing constant (see Supplementary Information). A priori, the variant allele frequency in a sample is exactly zero (*f* = 0) with some positive probability *c*_0_. The prior *π* is then of the following form

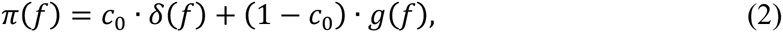

where *δ*(*f*) denotes the Dirac delta function and *g*(*f*) denotes a prior given the variant is present. We use a sample-specific prior function to account for the by multiple fold varying neoplastic cell content across samples (Supplementary Information; Fig. S2). The posterior probability that a variant is absent in a sample with low neoplastic cell content will be lower than in a sample with high neoplastic cell content despite the same *K* and *N* (Supplementary Information). The posterior probability that a variant is absent, denoted by *q*, and the probability that a variant is present, denoted by *p*, are

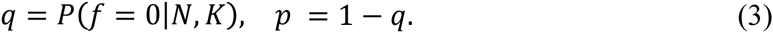

A variety of more sophisticated variant detection algorithms can be used here as long as the output can be converted to posterior probabilities of presence and absence. We calculate the probability of each mutation pattern for a particular variant by multiplying the corresponding posterior probabilities for each sample. Each mutation pattern has some positive probability, but those supported by the data are given much more weight.

A mutation pattern *v* is denoted as a binary vector of length |*S*| (total number of samples) where *v*_*s*_ is 1 if the variant is present in sample *s* and 0 if absent. The likelihood *L*_*μ*_(*v*) that a variant *μ* exhibits pattern *v* is

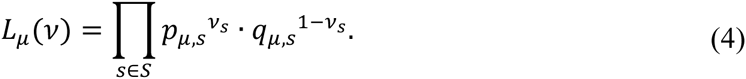

If the presence or absence of a variant in some samples is uncertain, the likelihood of any individual mutation pattern will generally be lower. The reliability score *ω*_*v*_ of each mutation pattern *v* (corresponding to a node in the evolutionary conflict graph; Fig. S3) is given by

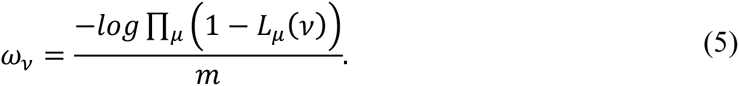

The argument of the logarithm denotes the probability that no mutation has pattern v and hence leverages the full sequencing information from all variants. With these scores (weights) normalized by the number of considered variants *m*, the minimum weight vertex cover of the evolutionary conflict graph corresponds to identifying the most reliable and evolutionarily compatible mutation patterns (see Supplementary Information for further details).

### Identifying reliable evolutionarily compatible mutation patterns

Given the calculated reliability scores, we efficiently find the most reliable and evolutionarily compatible mutation pattern for all variants via solving a Mixed Integer Linear Program^44^ (MILP). In the Supplementary Information we prove that finding these mutation patterns is equivalent to solving the *Minimum Vertex Cover problem*; one of Karp's original 21 *NP*-complete problems^39,55^. In the Minimum Vertex Cover problem one wants to find the minimum set of nodes in an undirected graph such that each edge in the graph is adjacent to one of the nodes in the minimum set. Therefore, by definition all edges are covered by the nodes in the minimum set. Similarly, we try to find the weighted set of nodes (here mutation patterns) with the minimal sum of reliability scores such that no evolutionary incompatibilities in the conflict graph remain. After this minimal set of nodes and their adjacent edges have been removed from the graph, we can easily infer an evolutionary tree since evolutionary conflicts no longer exist (i.e., all edges were covered and removed with the minimal set). The remaining set of mutation patterns is by definition the maximal set of evolutionarily compatible patterns (Supplementary Information).

In the evolutionary conflict graph *G* = (*V*, *E*), each node *i* ∈ *V* represents a different mutation pattern. For *n* samples, the number of nodes |*V*| is given by 2^*n*^. For each pair of evolutionarily incompatible mutation patterns *i* and *j*, there exists an edge (*i*, *j*) ∈ *E*. The weight (*c*_*i*_) of each node i is given by the reliability scores *ω*_*i*_ described in the Bayesian inference model section (Fig. S3).

The MILP to find the minimal-weighted set of evolutionarily incompatible mutation patterns is defined by the following objective function and constraints:

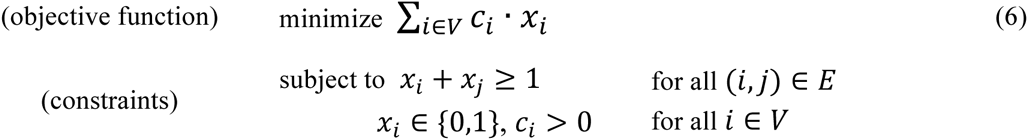

This formulation guarantees that the MILP solver finds the minimal value of the objective function such that all constraints are met and hence the nodes in the selected set cover all edges. The evolutionarily compatible and most reliable mutation patterns {*i* | *x*_*i*_ = 0} are given by the complement set of the optimal solution {*i* | *x*_*i*_ = 1} to the MILP.

### Inferring evolutionary trees

After the evolutionarily compatible mutation patterns {*i* | *x*_*i*_ = 1} have been identified and variants are assigned to their most likely evolutionarily compatible pattern based on the maximum likelihood weights given by the Bayesian inference model, the derivation of an evolutionary tree is a trivial computational task. In quadratic time (*O*(*n* · *m*)) of the input size we construct a unique phylogeny where *n* is the number of samples and *m* is the total number of distinct variants^56^. The branches where the individual variants are acquired follow from the inferred tree.

### Detecting subclones of distinct origin

Evolutionary incompatible mutation patterns with high reliability scores may indicate mixed subclones with distinct evolutionary trajectories (Fig. S9b, Fig. S10a). Recall that evolutionary incompatibility requires that the conflicting variants need to be present together in at least one sample. However, even if both variants are mutated in a statistically significant fraction in the same sample, these variants may not be present in the same cells and the evolutionary laws of an asexually evolving population may not be violated. If an evolutionarily incompatible mutation pattern exhibits a reliability score higher than expected from noise, Treeomics utilizes this evidence to infer subclones with distinct evolutionary trajectories and unidirectional spreading. A detailed pseudo-code is provided in the Supplementary Information. As outlined for prostate cancer case A22^10^, subsets (descendants) and supersets (ancestors) of the conflicting mutation pattern can simultaneously be identified and a comprehensive evolutionary tree is inferred (Fig. S9c). This approach also worked well among samples from the same tissue. After two subclones were separated in mixed samples from a prostate tumor^11^, 12643 (out of 12645) variants supported the inferred evolutionary tree (Fig. S10b). The remaining two variants were predicted to be false-positives by Treeomics. We performed extensive benchmarking of the subclone detection algorithm for various scenarios described in the following section (Fig. S12).

### *In silico* benchmarking

To assess the performance of Treeomics, we simulated metastatic progression according to a stochastic multi-type continuous-time branching process^48,57–59^ where metastases are seeded independently at random. Cells divide with birth rate *b* = 0.16, die with death rate *d* = 0.1555, and can leave the current site to successfully colonize a new site with probability *q* = 10^‒1148,60^. When a cell divides, it can accumulate a mutation with probability *u* = 0.045 (assuming a point mutation rate of 10^‒9^ and 45 megabases covered by Illumina exome sequencing^61^). The evolutionary process is initiated by a single advanced cancer that already accumulated driver gene mutations. Subsequently accumulated mutations, Single Nucleotide Variants (SNVs) and Copy Number Variants (CNVs), are assumed to be neutral^62,63^. Variants are acquired randomly across all chromosome pairs such that no two copy number events overlap (reasonable if the CNV mutation rate is low). SNVs and CNVs may overlap, in which case the timing of the events is used to determine the allele fraction of SNVs at the affected locus. CNV length is sampled from the observed length distribution in ref. 64. Cells in the primary tumor grow exponentially up to a fixed carrying capacity^65^. After *m* spatially-distinct metastases reached the detection size *M* = 10^7^, the simulation is stopped. Note that new metastases can also be seeded from previously seeded metastases.

To model the biopsy and sequencing process, a single sample consisting of one million cells of each of the *m* metastases consistent to the considered purity (30%, 45%, 60%, 75%, 90%) is subject to *in silico* sequencing. Metastases with a mixture of ancestries are simulated by random sampling from two distinct sites proportional to the tumor sizes at these sites. Sequencing depth is negative-binomially distributed with a given mean (50, 100, 200, 400, 800). A sequencing error rate of *e* = 0.5% is assumed. The simulation output is the number of variant and reference "reads" in each metastasis sample for each mutated locus present in at least 1% (or more for low sequencing depth) of any of the sampled metastases. An example for a simulated phylogeny is depicted in Fig. 3a. All the simulated phylogenies are available on github: https://github.com/johannesreiter/metastasesseeding.

We compared Treeomics to standard phylogenetic reconstruction (Maximum Parsimony^66^, Neighbor Joining^66^) and modern tumor phylogeny reconstruction methods (PhyloWGS^7^). Two different error metrics demonstrate the performance of Treeomics against existing methods: branching error and mutation matrix error score. The branching error quantifies the accuracy of the reconstructed coalescent relationships among distinct sites. From the true coalescent tree among metastatic sites, the collection of coalescent events among the sites is computed and compared to those predicted by the method. The branching error is defined as the fraction of true coalescent events missed by the reconstruction method. Since maximum parsimony and neighbor joining trees do not assume a perfect and persistent phylogeny, the branching error metric was used to compare these methods (Fig. 3, Fig. S11). The mutation matrix error score quantifies the accuracy of the reconstructed sequence of mutations acquired during an evolutionary process. For a tumor with *k* parsimony-informative mutations across *m* metastases, a *k* by *k* matrix *k* is constructed where *A*_*i,j*_ = 1 if mutation *i* is parental to mutation *j* and 0 otherwise. In PhyloWGS, where many phylogenies are sampled, this matrix is averaged over all samples. For a reconstructed phylogeny mutation matrix *Â*, the normalized error score is computed as Σ_*i,j*_(*A*_*i,j*_ – *Â*_*i,j*_)^2^/(*k*^2^ – *k*) Because PhyloWGS does not directly infer the coalescent relationship among sites, the mutation matrix error score was used in the benchmarking (Fig. 3, Fig. 12). Recall that only founder and parsimony-informative mutations were provided as input to PhyloWGS while Treeomics also had to deal with noisy private mutations. PhyloWGS was run with 2,500 MCMC iterations and 5,000 inner Metropolis-Hastings iterations for a maximum of 15 hours for each individual case. Increasing the number of samples and iterations did not significantly decrease the mutation matrix error score.

### Binary present/absent classification

We perform conventional binary present/absent classification of each variant to allow a comparison to the inferred classification used in our new approach. We scored each variant by calculating a p-value in all samples (one-tailed binomial test): 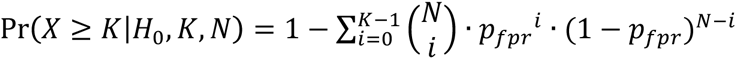 where *N* denotes the coverage, *K* denotes the number of variant reads observed at this position, and *X* denotes the random number of false-positives. As null hypothesis *H*_*0*_, we assume that the variant is absent. Similar to Gundem et al.^10^, we assumed a false-positive rate (*p*_*fpr*_) of 0.5% for the Illumina chip-based targeted deep sequencing. We used the step-up method^67^ to control for an average false discovery rate (FDR) of 5% in the combined set of *p*-values from all samples of a patient. Variants with a rejected null hypothesis were classified as present. The remaining variants were classified as absent.

### Code availability

The source code and a manual for Treeomics, as well as multiple examples illustrating its usage, are provided at https://github.com/johannesreiter/treeomics. Treeomics v1.3 was used for the entire analysis. The tool is implemented in Python 3.4. The inputs to the tool are the called variants and the corresponding sequencing data, either in tab-separated-values format or as matched tumor-normal VCF files. As output, Treeomics produces a comprehensive HTML report (Supplementary File 1) including statistical analysis of the data, a mutation table plot and a list of putative artifacts (false-positives, well-powered and under-powered false-negatives). Additionally, Treeomics produces evolutionary trees in LaTeX/TikZ format for high-resolution plots in PDF format. If circos^68^ is installed, Treeomics automatically creates the evolutionary conflict graph and adds it to the HTML report. Treeomics also supports various filtering (e.g., minimal sample median coverage, false-positive rate, false-discovery rate) for an extensive analysis of the sequencing data. Detailed instructions for the filtering and analysis are provided in the readme file in the online repository. For solving the MILP, Treeomics makes use of the common CPLEX solver (v12.6) from IBM.

## ACKNOLEDGEMENTS

We thank Martin Chmelik, Alison Hill and Adeeti Ullal for valuable discussions. This work was supported by the European Research Council (ERC) start grant 279307: Graph Games (J.G.R., C.K.), Austrian Science Fund (FWF) grant no P23499-N23 (J.G.R., C.K.), FWF NFN grant no S11407-N23 RiSE/SHiNE (J.G.R., C.K.), a Landry Cancer Biology Fellowship (J.M.G.), National Institutes of Health grants CA179991 (C.I.-D.), F31CA180682 (A.M.-M.), CA43460 (B.V.), the Lustgarten Foundation for Pancreatic Cancer Research, the The Sol Goldman Center for Pancreatic Cancer Research, the The Virginia and D.K. Ludwig Fund for Cancer Research, the John Templeton Foundation and a grant from B. Wu and Eric Larson. We thank Bashashati et al. (2013), Cooper et al. (2015), and Gundem et al. (2015) for sharing their comprehensive data sets. Benchmarking was performed on the Odyssey cluster supported by the FAS Division of Science, Research Computing Group at Harvard University.

## SUPPLEMENTARY INFORMATION

1) Supplementary Information: Figures S1-S13, purity analysis, Bayesian inference model, subclone detection algorithm, mathematical proofs, Treeomics manual
2) Supplementary Tables S1-S4
3) Supplementary File S1: Example of an HTML analysis report produced by Treeomics
4) Source code and examples: https://github.com/johannesreiter/treeomics *(released after publication)*
5) Simulated sequencing data: https://github.com/johannesreiter/metastasesseeding (>1.5GB of data, *released after publication*)

